# Single photon kilohertz frame rate imaging of neural activity

**DOI:** 10.1101/2022.05.23.493031

**Authors:** Tian Tian, Yifang Yuan, Srinjoy Mitra, Istvan Gyongy, Matthew F Nolan

## Abstract

Establishing the biological basis of cognition and its disorders will require high precision spatiotemporal measurements of neural activity. Recently developed genetically encoded voltage indicators (GEVIs) report both spiking and subthreshold activity of identified neurons. However, maximally capitalising on the potential of GEVIs will require imaging at the millisecond time scales, which remains challenging with standard camera systems. Here we report application of single photon avalanche diode (SPAD) sensors to imaging neural activity at kilohertz frame rates. SPADs are electronic devices that when activated by a single photon cause an avalanche of electrons and a large electric current. We use an array of SPAD sensors to image individual neurons expressing genetically encoded voltage indicators. We show that subthreshold and spiking activity can be resolved with shot noise limited signals at frame rates of up to 10 kHz. SPAD imaging was able to reveal millisecond scale synchronisation of neural activity in an ex-vivo seizure model. SPAD sensors may have widespread applications for investigation of millisecond timescale neural dynamics.

**Table of contents:** The high temporal precision of single photon avalanche diodes (SPADs) is leveraged to record neural activity reported by genetically encoded voltage indicators. Sub-threshold and spiking activity of single neurons was resolved with shot noise limited signals at frame rates of up to 10 kHz. SPAD sensors may have widespread applications for neural imaging at high frame rates.

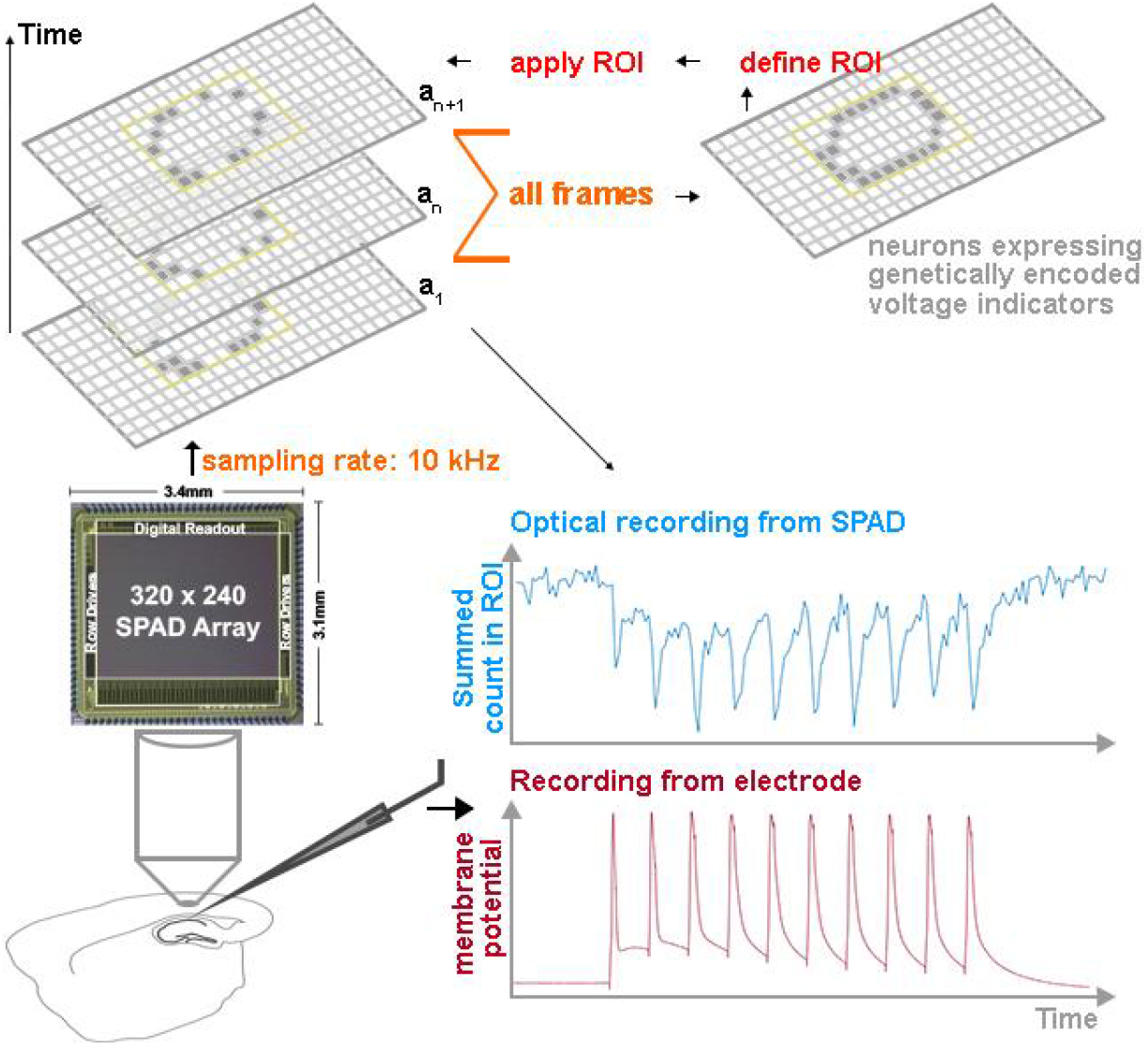

## Introduction

A long-standing goal of neuroscience is to resolve the activity of ensembles of identified neurons with high spatiotemporal precision. Genetically encoded voltage indicators (GEVIs) can reveal subthreshold activity and resolve spike timing with millisecond resolution from identified neuronal populations^1–3^. These are substantial advantages over fluorescent genetically encoded calcium indicators^4,5^. However, fluorescence imaging of neural activity reported by GEVIs is challenging due to their millisecond dynamics and limited photon budget^6^. An ideal imaging system would provide efficient shot-noise limited quantification of GEVI fluorescence at frame-rates above the Nyquist limit for sampling action potentials, which typically have durations on the order of a millisecond.

To date, electron multiplying CCDs (emCCD) and scientific CMOS (sCMOS) cameras have typically been employed to image GEVIs (Supplementary Table 1)^1–3,7^. These sensors face constraints in achieving high frame rates while minimising the impact of inherent noise sources. Higher frame-rates are a particular challenge as they demand a faster operation of the read-out electronics, which can conceal true signals through increased electronic readout noise^8^. State-of-the-art emCCD cameras have low effective read noise and can achieve single-photon sensitivity^8^. However, the limited frame-rate of the emCCD cameras (~100 frames per second (fps)) does not allow kilohertz frame-rate imaging and the amplification process also introduces noise that effectively lowers the signal-to-noise ratio and their quantum efficiency^9^. State-of-the-art sCMOS suffer from inherent dark noise and read noise^8^, which makes it difficult at high frame rates to separate the weak signal generated by the incidence of a limited number of photons from spurious noise events.

The single photon avalanche diode (SPAD) is a photodiode that is reverse biased above its breakdown voltage, so that a single photon incidence at its photosensitive region creates an electron-hole pair that triggers an avalanche of secondary carriers and a large electric current^10,11^. A SPAD is thus capable of single photon detection with the rising edge of the voltage pulse encoding the time of arrival of the photon. Local (in-pixel) circuitry then actively lowers the SPAD bias voltage to below the breakdown voltage to stop the avalanche and subsequently recharge the bias voltage to its initial value above the breakdown voltage, restoring the SPAD’s sensitivity for the next photon detection event. This cycle typically takes on the order of tens of nanoseconds, enabling SPADs to perform ultra-high frame-rate time-resolved imaging such as capturing light-in-flight^12^. Because SPADs can detect the time at which a photon hits the sensor, their effective frame-rates are set by temporal binning and can be arbitrarily chosen after data acquisition according to the specific experimental demands so as to capture fast optical dynamics and maximise the available photon budget. SPAD-based image sensors have attained deep sub-electron read noise (0.06 e^-^ to 0.17 e^-^, SPAD sensor used in this study^13^). Therefore, at high frame rates the SPAD read noise is effectively negligible and images can be obtained at the shot-noise limit^13^. As a result, in the low photon regimes encountered in high-speed imaging, the signal-to-noise ratio of a SPAD can exceed that of an sCMOS^14^. Furthermore, in contrast to conventional emCCD and sCMOS cameras, temporal and/or spatial oversampling and binning can be carried out without any noise penalty^15^. Although the application of SPADs to biological imaging has in the past been constrained by small array sizes, as well as large pixels with low fill factor, technological advances in these areas are making biological applications increasingly feasible^11^.

Here, we evaluate the suitability of SPAD-based imaging for recording neuronal activity reported by GEVIs. We show that SPAD image sensors can resolve individual neuronal sub-threshold and spiking activity reported with the GEVI Voltron-JF525-HTL with shot noise limited signals at frame rates of up to 10 kHz. They can also reveal spiking activities of individual neurons in neural ensembles during seizure-like events induced by 4-AP. SPAD sensors may have widespread applications for neural imaging at high frame rates.

## Results

We utilised a 320 × 240 SPAD array image sensor (SPCImager, Supplementary Table 1) with 8 μm pixel pitch and 26.8% fill factor (FF) and a peak photon detection probability of 35% at 450 nm^13^ (Figure 1A). When operating in binary mode each SPAD pixel produces a time-domain sequence of 0 s (no photon detected) or 1 s (at least one photon detected) and the raw output of the SPAD at each exposure is the summed binary pixels in space, also known as a “bit plane”. A rolling shutter was used to enable back-to-back exposure at the maximum frame-rate (close to 10 kHz), so that the exposure time for each bit-plane is around 100 μs. The SPAD sensor was paired with an FPGA board (Opal Kelly XEM6310) that controls the acquisition of image data, relaying a continuous stream of bit-planes to a PC over a USB 3.0 link^15^. Figure 1B shows a schematic of the experiment setup of the SPAD sensor incorporated with a microscope and electrophysiological recording set up. Figure 1C shows the temporally oversampled greyscale image of a neuron expressing the GEVI Voltron-JF525-HTL, obtained by binning 10,000 consecutive bit-planes over a 1-second long recording^13,16^.

**Figure 1.**
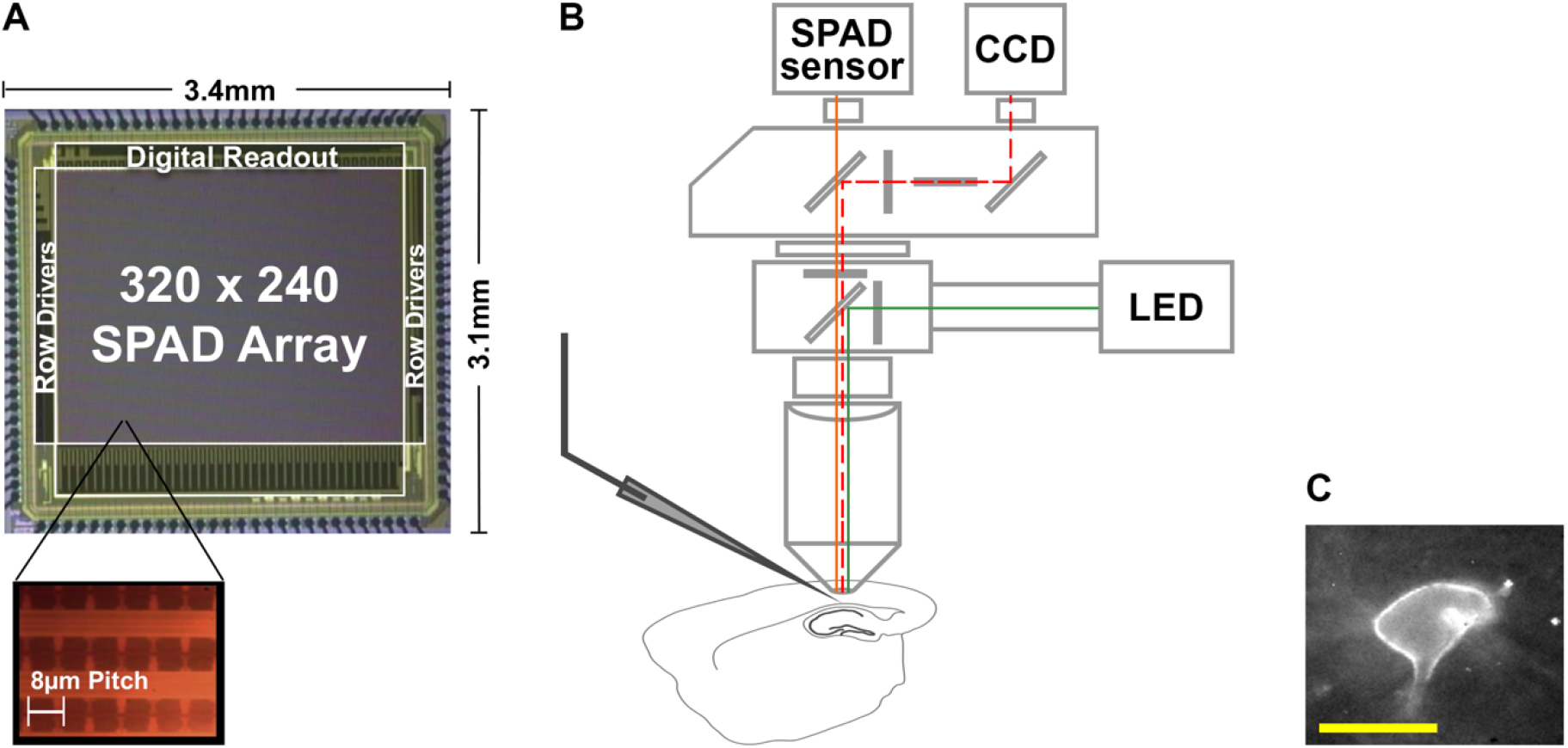
Photomicrograph of the SPAD image sensor and schematic representation of experiment setup. (A) Photomicrograph of the SPAD image sensor and magnified image of the pixel array. (B) Schematic of the experiment setup with coloured lines indicating the simplified light path. The CCD camera was used for localisation of neurons and positioning patch-clamp electrodes. Optical dynamics of the Voltron-JF525-HTL were recorded by the SPAD sensor. (C) An example image captured by the SPAD array of a patch-clamped neuron expressing Voltron-JF525-HTL. The Voltron is mainly concentrated on the membrane of the soma. The distortion of the membrane caused by the pipette results in a local increase in fluorescence. Scale bar for neuron image: 10μm.

To test the suitability of the SPAD sensor for fluorescence voltage imaging, we used a viral approach to transduce cortical and hippocampal neurons with the GEVI Voltron-JF525-HTL^1^ and made whole-cell patch clamp recording from GEVI-expressing neurons in ex vivo hippocampal and neocortical brain slices. We used the patch-clamp electrode to manipulate the membrane potential of the recorded neurons and simultaneously recorded the fluorescence changes of Voltron-JF525-HTL using the SPAD sensor. In sparsely labelled preparations^17^, we observe fluorescence concentrated to the membrane of the soma of neurons expressing the soma-targeted Voltron-JF525-HTL (Figure 1C, Figure 2A). For post-processing, the region of interest (ROI) was selected manually and a binary mask was applied to the ROI, allowing isolation of the fluorescence-positive pixels within the ROI and from which we obtained the time-dependent traces from each neuron (Figure 2). We took advantage of the negligible read noise and temporal oversampling of the SPAD sensor and adjusted the effective frame-rate by temporally binning consecutive bit-planes to optimise detection of sub-threshold voltage changes or supra-threshold spikes (Figure 2, Supplementary Figure 1). Temporal binning was carried out by generating an average of summed photon count within the ROI of consecutive bit-planes to reach an effective frame-rate suitable for the specific biological event (Figure 2B-C).

**Figure 2.**
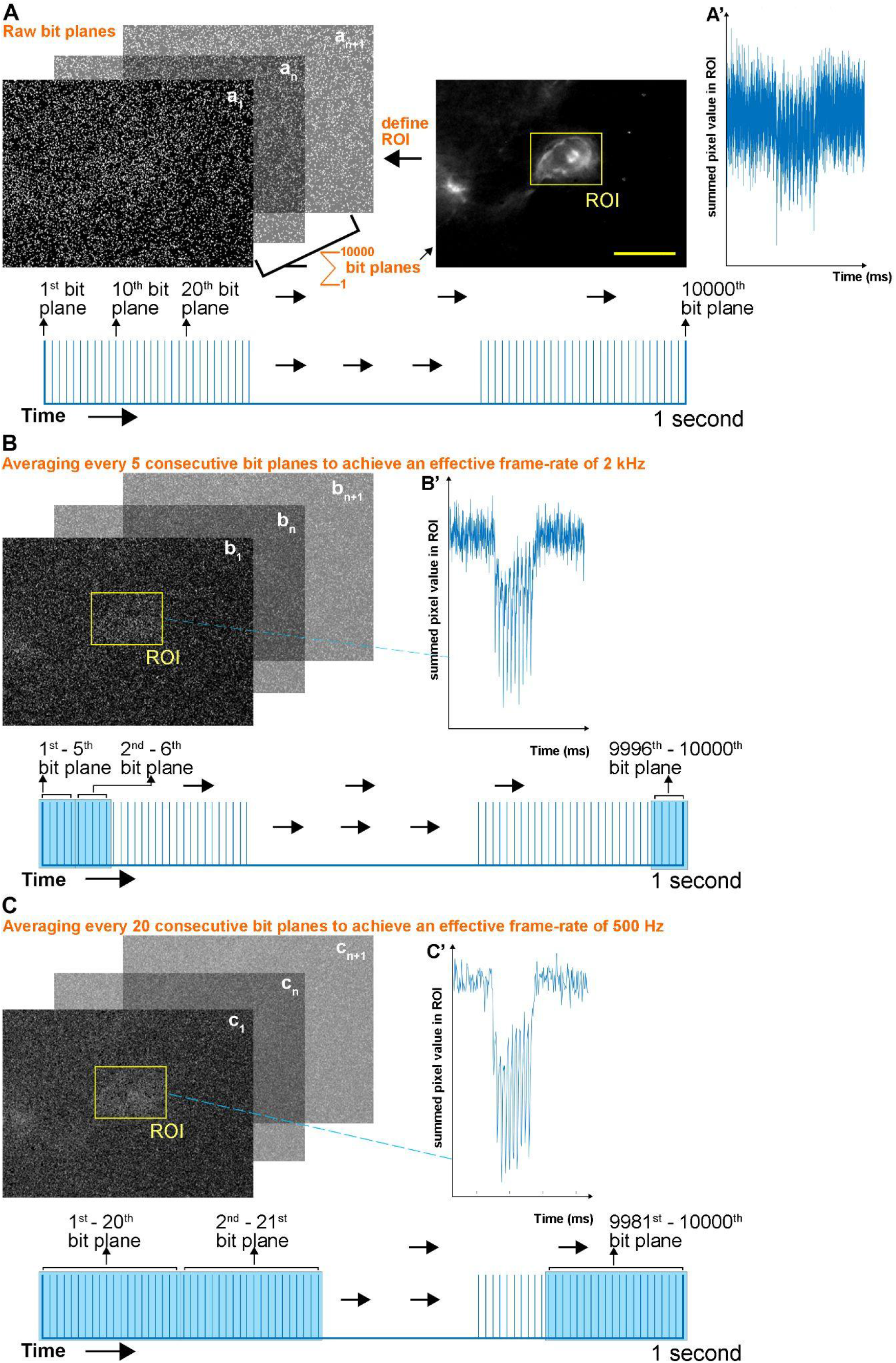
Strategy for temporal binning of bit planes. (A) To define ROIs, 10000 raw bit planes were collected from a 1-second-long recording and then summed to yield an image of the field of view. The summed photon counts within the ROI of each raw bit plane are plotted in (A’) as a function of time during delivery of 10 spikes at 100 Hz. (B) To perform temporal binning a set number of consecutive bit planes were averaged (in this example 5 consecutive bit planes were averaged, therefore achieving an effective frame rate of 2 kHz). Summed photon counts within the ROI of the averaged bit planes are plotted in B’ as a function of time (corresponding to the unbinned data in A’). The images in the top left show the field of view after temporal binning of 5 consecutive bit planes was applied. (C) As for (B) with temporal binning of 20 consecutive bit planes to achieve an effective frame rate of 500 Hz. Scale bar: 10 μm.

The SPAD image sensor captured slow changes (tens of milliseconds) in baseline membrane potentials reported with Voltron-JF525-HTL in acute brain slices, detecting subthreshold hyperpolarisation and depolarisation of the membrane potential in response to current and voltage stimuli (Figure 3 and Supplementary Figure 2). Temporally binning more consecutive frames of the optical traces of subthreshold events significantly increased the SNR, but not the ΔF/F at each current or voltage step (Supplementary figure 1A-F). Based on these considerations we chose an effective frame-rate of 100 Hz for analysis of subthreshold activity. We found that steady-state changes in baseline membrane potential (ΔV) were strongly correlated with changes in ΔF/F in both current clamp (R^2^ = 0.990 ± 0.00200, p < 0.0001, Figure 3E) and voltage clamp (R^2^ = 0.976 ± 0.00355, p < 0.0001, Supplementary Figure 2C-E) and the relationship between ΔV and ΔF/F appeared linear. The kinetics of membrane potential responses to current steps were also captured by the SPAD sensor (Figure 3C-D), with no significant difference in the rise and decay time constant calculated from electrophysiological and optical recordings (p = 0.511, χ(1)^2^ = 0.4316, likelihood ratio test, n = 9 cells, Figure 3F).

**Figure 3.**
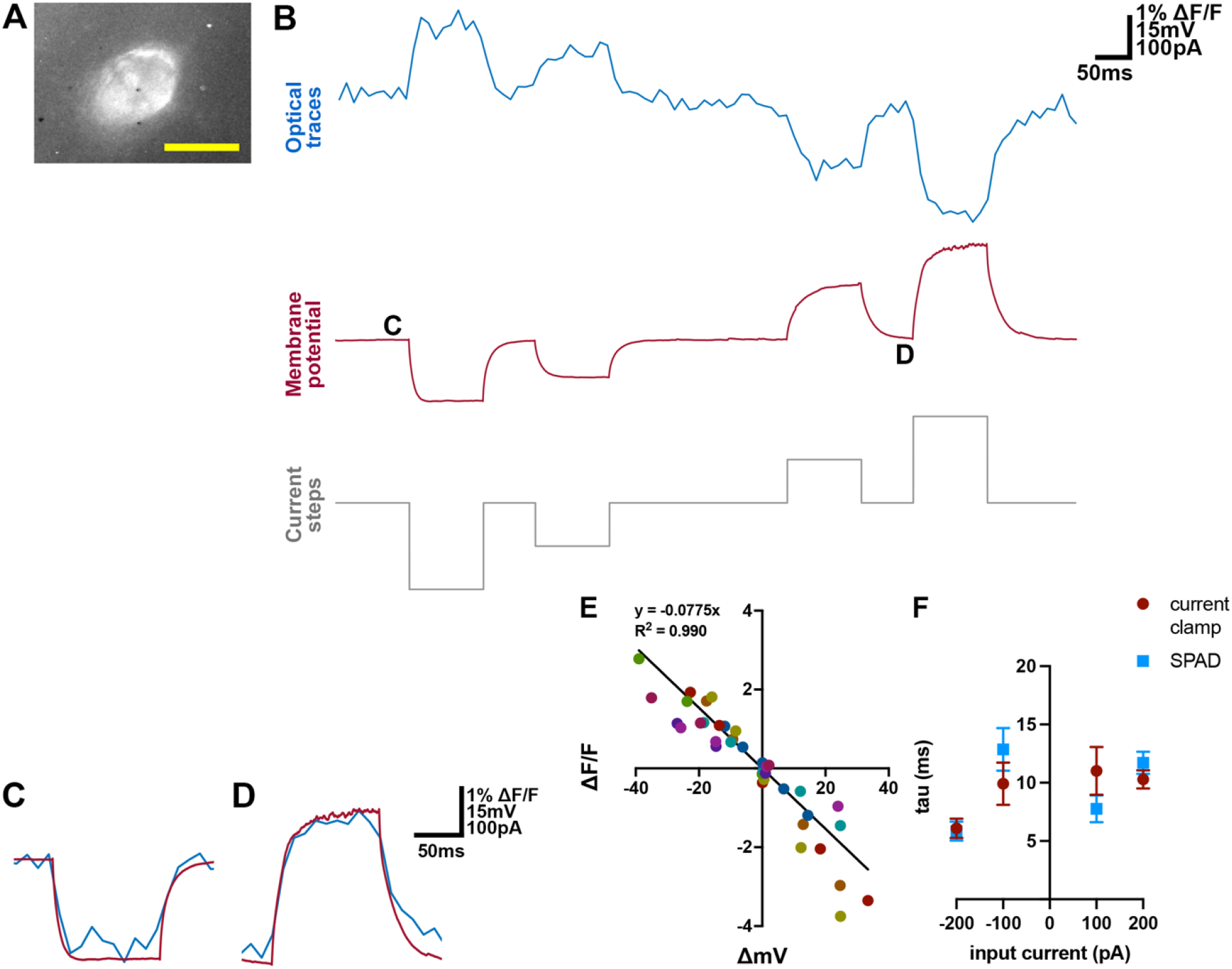
Detection of subthreshold membrane potential changes in the soma reported with Voltron-JF525-HTL. (A) Image captured by the SPAD array of a patch-clamped neuron expressing Voltron-JF525-HTL, from which the traces in (B-D) was recorded. Scale bar for neuron image: 10μm. (B) Simultaneous optical (blue) and electrical (red) recordings of membrane potential changes in response to current steps (grey). The optical traces were captured by the SPAD sensor at a sampling rate of 9.9384 kHz, low pass filtered at 2 kHz and temporally binned at 100 Hz. The electrical traces were sampled at 20 kHz. (C-D) Segments from B on an expanded time base with the optical (blue) and electrical (red) traces superimposed. (E) Change in Voltron signal (ΔF/F) as a function of membrane potential change (ΔmV) in response to various current inputs (n = 9 cells). Data from each cell is marked with a different colour. The line indicates the fit obtained with linear regression with intercept set at x = 0 and y = 0 (p < 0.0001). (F) Time constants (mean ± SEM) estimated from fitting responses to current steps plotted as a function of the current step amplitude. There was no significant difference between time constants estimated with electrical and optical methods (p = 0.511, χ(1)^2^ = 0.4316, likelihood ratio test, n = 9 cells,).

We next tested if the SPAD sensor can capture individual action potentials. Although ΔF/F of spiking events decreased with temporal binning, the SNR increased significantly at lower effective frame rates (Supplementary figure 1G-L). To capture the fast optical transient of Voltron during an action potential, we temporally binned 10 consecutive SPAD bit-planes to reach an effective frame rate of 1 kHz. As shown in Figure 4A-D, the optical traces recorded by the SPAD sensor tracks single spikes in action potential trains evoked by 25 Hz and 100 Hz current pulses (See also Supplementary figure 3). The relative fluorescence changes, quantified either by the SNR or the ΔF/F of single spikes, were comparable to the original report of Voltron-JF525-HTL (Figure 4E-F)^1^. Complex spikes evoked by a single pulse of current injection were also reliably reported by the SPAD signal (Supplementary figure 3B). Occasionally, the expression of the soma-targeted Voltron-JF525-HTL can also be seen in subcellular structures close to the soma (Figure 4G). In these cases, the SPAD signal reported current-evoked action potential trains in both the soma and in the subcellular structure (Figure 4H), with reduced peak amplitude in the subcellular structure (Figure 4I).

**Figure 4.**
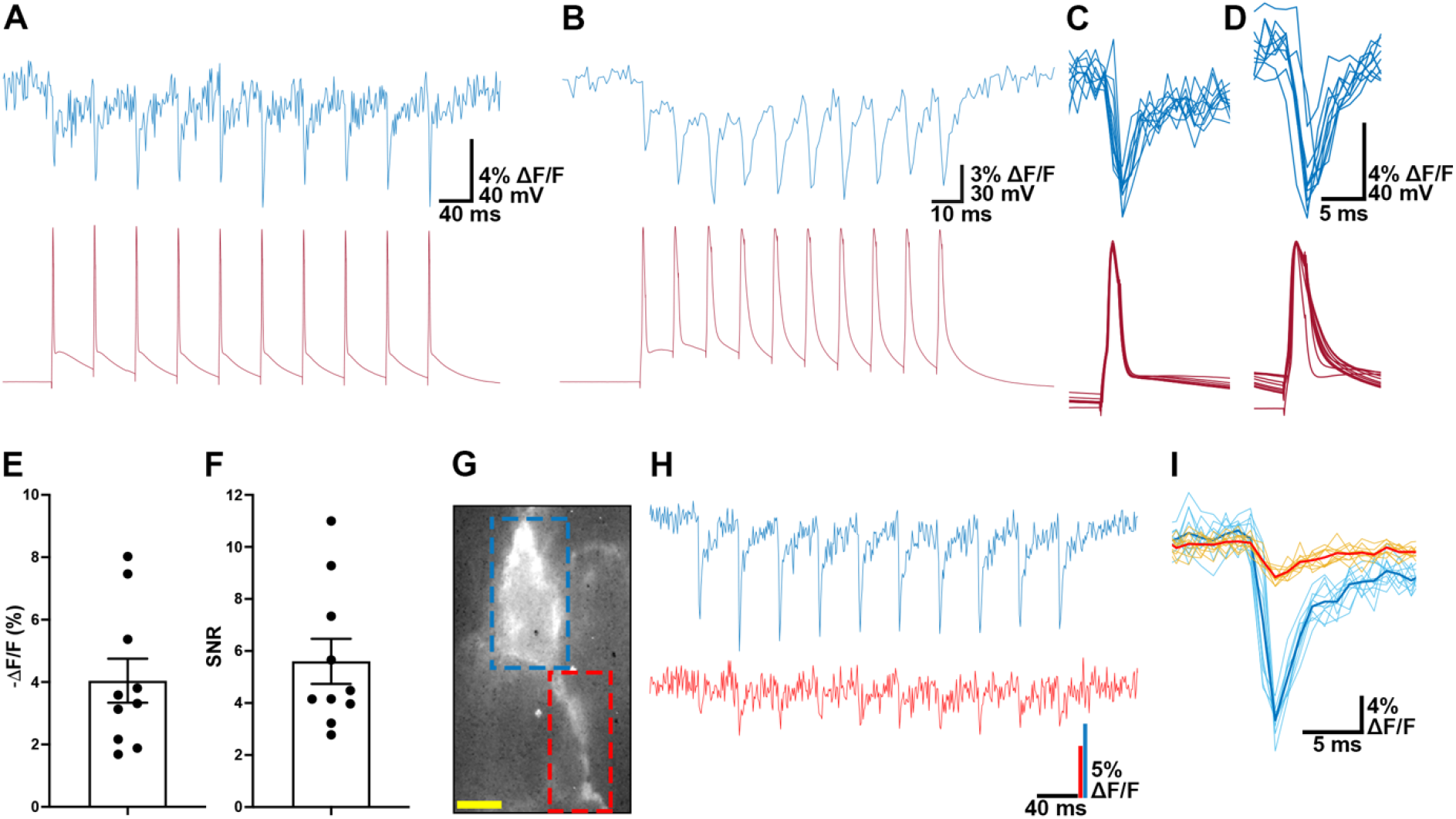
Detection of action potentials in the soma and subcellular structures reported with Voltron-JF525-HTL. (A-B) Simultaneous optical (blue) and electrical (red) recordings of action potentials in response to 10 current pulses at 25Hz (A) and 100Hz (B). Pulse amplitude is 2 nA duration is 2 ms. The optical traces were originally captured by the SPAD image sensor at a sampling rate of 9.9384 kHz, then low pass filtered at 2 kHz and temporally binned at 2 kHz. The electrical traces were sampled at 20 kHz. (C-D) Action potentials from (A-B) plotted on an expanded time base. (E-F) Mean (± SEM) ΔF/F (E) and SNR (F) of peak fluorescence response in ex vivo brain slices. Each neuron fired 10 APs at 25Hz, the output of each cell was the mean over these 10 spikes (n = 10 cells). SNR was calculated as ΔF / (standard deviation of baseline fluorescence F). (G-I) Imaging fluorescence changes in a primary dendrite (blue square in G) as well as the adjacent soma (ROI shown by the red square in G) detects action potentials in both areas in responses to a train of current pulses (H). The amplitude of the action potential is smaller in the dendrite (I) (solid lines indicating average spike waveforms are overlaid on the individual responses). Scale bar for neuron image: 10μm.

We next tested if the SPAD sensor can report activities of individual neurons in neural ensembles during epileptiform activity. We induced seizure-like events in hippocampal slices by application of 4-aminopyridine (4-AP). Interneurons in the striatum oriens layer of CA1 fired bursts of action potentials that were detected similarly with patch-clamp recordings and simultaneously with SPAD imaging of Voltron-JF525-HTL signals (Figure 5A-B, Supplementary Figure 5). The SPAD sensor reported action potentials and subthreshold events from multiple neurons during low frequency and bursting activity (Figure 5B-D, Supplementary figure 5A-B, D-E). Peristimulus time histogram (PTSH) analysis of spiking activities of cell pairs showed evidence of millisecond scale synchronisation (Figure 5E) and coordinated firing with lags of up to 100 ms (Figure 5F). Consistent with this, cell pairs showed strong sub-threshold signal correlations reflecting synchronous depolarisations during epileptiform bursts of activity (Figure 5B, G).

**Figure 5.**
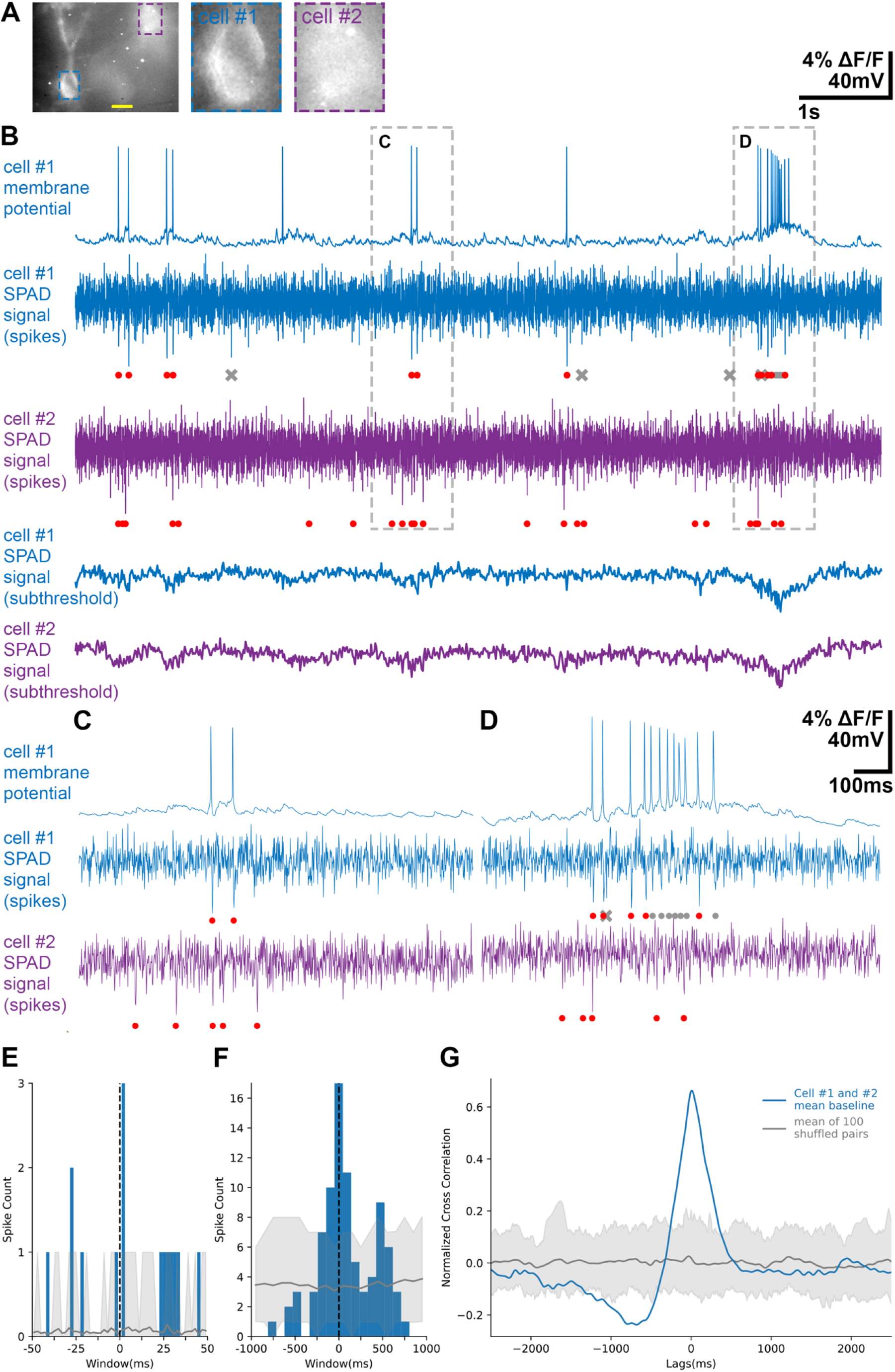
Coordination of spiking activity during seizure-like events. (A) A field of view of recorded neurons in the striatum oriens layer of CA1 expressing Voltron-JF525-HTL, taken with a SPAD image sensor using a x20 water immersion objective. Scale bar: 10 μm. (B) Activity of the neurons in (A). The membrane potential changes of Cell #1 recorded with the parch-clamp electrode is shown above the optical traces captured by the SPAD image sensor for Cell1 #1 (blue) and Cell #2 (purple) and processed to isolate spikes and sub-threshold activity (see Supplementary Figure 5). (C-D) Dotted-line areas in (B) are expanded, showing low frequency firing (C) and bursts (D). When the optical trace of cell #1 was compared with the ground-truth electrical trace, the correctly detected spikes were marked with a red dot, the false negative spikes were marked with a grey dot, the false positive spikes were marked with a grey cross. (E-F) Peristimulus time histogram (PSTH) of spiking activity between Cell #1 and Cell #2 from (B). With the spikes of Cell #1 being set as reference events, the sum of spike counts of Cell #2 was plotted with 2 ms bins (E), and the sum of spike counts of Cell #2 was plotted with 100ms bins (F). Blue trace: PSTH of the two traces from the cell pair. Grey line: mean PSTH of shuffled data. Grey shaded area: 5%-95% percentile of shuffled data. 100 randomly shuffled cell pairs were generated by shuffling the spike times of the original two traces. (G) Cross-correlogram of subthreshold signals between the cells in (B). Blue trace: mean normalized cross correlation of ten 10-second-long optical recordings of subthreshold baseline fluctuations. Grey line: mean normalized cross correlation of shuffled data. Grey shaded area: 5%-95% percentile of shuffled data. 100 randomly shuffled cell pairs were generated by pairing 10-seconds recording traces from different cells in different brain slices.

## Discussion

We demonstrate kilohertz frame-rate voltage imaging with subcellular resolution in ex vivo mouse brain slices using a SPAD imaging sensor. We recorded both supra- and subthreshold activity of neurons during electrically triggered responses and seizure-like events induced by 4-AP. The SPAD imaging sensors we use here can readily be applied to existing microscopes hence giving access to voltage imaging in most in vitro, ex vivo and head fixed settings.

The ability of SPADs to detect the time of photon arrival makes them well suited to imaging fast changes in neuronal membrane potential. The effective frame rate can be chosen arbitrarily in post-processing; higher frame-rates can be achieved by shorter binning intervals, although at the cost of decreased SNR. Given their low intrinsic noise at high frame rates the quality of the imaging signal is essentially shot noise limited. Our existing SPAD imaging sensor can readily be applied to existing microscopes, hence giving access to voltage imaging in most in vitro, ex vivo and head fixed settings. SPAD-based sensors also have the advantage that their operation does not require bulky cooling. This may make them well suited to development of miniaturised imaging systems, for example to monitor neural activity in freely moving animals^18^.

The primary constraints for imaging neuronal membrane potential via SPAD sensors come from their quantum efficiency and from the properties of the GEVIs used. While our experiments establish that a SPAD sensor with an external quantum efficiency of ~10% is sufficient for imaging activity reported by GEVIs, recently developed SPADs^19^ featuring multibit pixels and efficiency > 50% will likely lead to substantial additional improvements in signal to noise ratios, with greater proportion of detected photons reducing the shot noise. Further improvement in brightness and sensitivity of voltage indicators^20–22^ will further enhance detection of voltage signals in subcellular compartments and more densely-labelled samples.

## Methods

### Animals

All animal experiments were conducted in accordance with the UK Animal (Scientific Procedures) Act 1986 and were approved by University of Edinburgh’s Animal Ethics Committee.

### Injection of viruses and dye

Male and female wild-type C57BL/6Crl mice aged between 21 and 40 days were anesthetized with isoflurane, mounted in a stereotaxic frame, and a small craniotomy made above the target region. To achieve sparse labelling, a mixture of pENN-AAV-hSyn-Cre-WPRE-hGH (Addgene # 105553-AAV1) (titer ≥ 2×10^9^ vg/mL) and pAAV-hsyn-flex-Voltron-ST (Addgene # 119036-AAV1) (titer ≥ 1×10^12^ vg/mL) injected into the primary visual cortex (stereotaxic coordinates AP −3.8, ML 3.0, DV −0.3 and −0.6) and CA1 (AP −3.8, ML 3.0, DV −1.3). Injection coordinates were calculated relative to bregma and 200 nl of the virus mixture was injected at each coordinate. JF525-HTL (100 nMol, Lavis Lab, Janelia Research Campus, HHMI) was mixed with 20μl of DMSO (Sigma, D2650-5X5ML), 20μl of Pluronic™ F-127 (20% Solution in DMSO) (ThermoFischer Scientific P3000MP) and 80 μl of 1x PBS and administered intravenously into the lateral tail vein of mice 24-72 hours prior to experimentation.

### Electrophysiology and fluorescence imaging in mouse brain slices

Preparation of brain slices and electrophysiological recordings were carried out as described previously ^26^. Briefly, sagittal brain slices were prepared from male and female wild-type C57BL/6Crl mice aged between 60 and 100 days. Mice were sacrificed by cervical dislocation. The brains were quickly removed and placed in ice-cold (2-4°C) cutting artificial cerebrospinal fluid (ACSF) (pH 7.4) containing (in mM): 86 NaCl, 1.2 NaH2PO4, 2.5 KCl, 0.5 CaCl2 (1M), 7 MgCl2 (1M), 25 NaHCO3, 25 Glucose, 50 Sucrose, and aerated with 95% O2, 5% CO2. The injected hemispheres were mounted on a Vibratome (Leica VT 1200, Leica Microsystems) and cut at 400 μm thickness, then transferred to standard ACSF containing (in mM) 124 NaCl, 1.2 NaH2PO4, 2.5 KCl, 2 CaCl2, 1 MgCl2, 25 NaHCO3, 20 Glucose and incubated at 37°C for 15 minutes. The slices were stored at room temperature in a submerged chamber constantly aerated with 95% O2, 5% CO2 for at least 1 hr before being transferred to the recording chamber perfused with standard ACSF at a flow rate of 3 ml/min at 33°C

The brain slices were first visualised with a digital CCD camera (SciCam Pro, Scientifica) mounted on an upright microscope (BX51-WI, Olympus) using either a 40x water-immersion objective lens (1.0 N.A., LUMPLFLN 40XW Olympus) or a 20x water-immersion objective lens (1.0 N.A., XLUMPLFLN 20XW, Olympus). For epifluorescent imaging of JF525 dye, a green LED (Thorlabs M530L4) was used with a dichroic filter set (Semrock FF520-Di02, FF01-500/24, and FF01-562/40). Recording pipettes were pulled from borosilicate capillary glass (Havard Apparatus, 30-0060) on a horizontal electrode puller (P-97, Sutter Instruments) to a tip resistance of 4-6 MΩ and filled with K-gluconate-based internal solution (in mM: 130 K Gluconate, 10 KCl, 10 HEPES, 2 MgCl2 (1M), 0.1 EGTA, 2 Na2ATP, 0.3 Na2GTP, 10 NaPhosophoCreatine, Biocytin 0.5%, pH=7.0-7.5, 290-300mOsm). Recording pipettes were positioned with a micromanipulator (Sensapex). Patch-clamp recordings were performed with a Multiclamp 700B amplifier (Molecular Devices), with membrane potential sampled at 20 KHz, filtered at 10 KHz with the built-in 4-pole Bessel Filter, and digitised (National Instrument, USB-6212). The command waveforms of electrical stimuli were generated, and the electrophysiological recordings were acquired using Axograph 1.7.6. Axograph was also used to generate signals to synchronise electrophysiological recordings and the SPAD imaging.

When whole-cell configuration was achieved, the light path of the microscope was switched to focus into the SPAD image sensor. To record baseline membrane potential changes, stimulation protocols of current steps (100 ms baseline, −200 pA to 200 pA, 100 ms per step, 100 pA increment) in current clamp and voltage steps (100 ms baseline, −50 mV to 30 mV, 100 ms per step, 20 mV increment) in voltage clamp were applied. In voltage clamp, cells were held at −70 mV. To record action potentials, a train of 10 current pulses (2 nA, 2 ms) was given to the neurons, with the frequency ranging from 25 Hz to 100 Hz.

To record neural activity during seizure-like events induced by 4-AP, one electrode was placed to record field potential whereas another electrode was used to achieve whole-cell configuration in a Voltron labelled neuron. When the whole-cell configuration was achieved, 200 μM 4-AP was added to the ACSF to induce seizure-like activity. When changes in field potential were recorded, indicating the onset of the seizure-like events, the SPAD imaging sensor was then started to record for 100 seconds (10 consecutive epochs of 10-second-long recordings).

### SPAD imaging sensor

The sensor chip was fabricated in STMicroelectronics 130nm imaging CMOS technology^13^. A detailed description can be found in ^13^. We developed custom FPGA firmware to allow the camera to be triggered directly from the electrophysiology software (Axograph 1.7.6) via the digitiser board (National Instrument, USB-6212). In all experiments raw bit frames were captured from the SPAD image sensor at a sampling rate of 9.9384 kHz.

### Data analysis and statistics

For composing images, postprocessing algorithms were used to correct both the dark count rate of the pixels and the logarithmic response of the pixels^27^. Regions of interest (ROI) were selected manually and a binary mask was applied to allow isolation of the fluorescence-positive pixels. For each bit plane counts were summed within the ROI and values for all bit planes concatenated to produce an optical trace. When generating time traces, only logarithmic response of the pixels were corrected^26^. For each optical trace, high-frequency noise was removed by applying a low-pass filter at 2 kHz. Temporal binning was carried out by calculating the mean value of every consecutive bit planes (Python 3.8) to obtain a smoothed trace (e.g. binning 10 consecutive planes gives an effective sampling rate of 1kHz). For investigation of responses to subthreshold current steps and voltage steps the mean fluorescence intensity of the first 100 ms within the ROI was used as the baseline fluorescence (F). For the action potential dataset, each spike was identified using the data processing pipeline provided in ^1^, and the peak intensity was read accordingly. ΔF/F was calculated as the peak intensity of each spike minus the baseline fluorescence of the same optical trace, divided by the baseline fluorescence. SNR of each spike was calculated as the peak intensity of each spike minus the baseline fluorescence of the same optical trace, divided by the standard deviation of the baseline fluorescence. The ΔF/F and SNR of each trace was the mean of ΔF/F and SNR of all the spikes identified in that optical trace. For the subthreshold activity (current step and voltage step) dataset, the ΔF was calculated as the mean fluorescence at the following time intervals of the optical trace: 120 to 180 ms, 290 to 350 ms, 460 to 520 ms (voltage step only), 630 to 690 ms, 800 to 860 ms minus the baseline fluorescence. The ΔF/F and SNR of each step was calculated as these values divided by the baseline fluorescence or the standard deviation of the baseline fluorescence. For each trace, simple linear regression was applied with intercept set at x = 0 and y = 0 (Excel Version 16.54). To calculate the time constant (tau) of each current step, the upward or downward deflection of 100 ms of electrophysiology and optical recordings after the onset of each current step were fitted with the nonlinear regression (curve fit) function in Prism (Version 9.1.2). The constant of the best-fitted nonlinear regression equation was the tau. Linear mixed effect models (LMEs) were fitted using lme 4 1.1-12^28^ in R v.4.1.2 (R Core Team, 2014).

The optical traces from the 4-AP experiments were first low-pass filtered at 2kHz to remove high-frequency noise. Subthreshold activities was revealed by temporally binning the low-pass filtered raw traces at 100 Hz and baseline drift was removed by applying the detrend command (no Type specified) in MATLAB (R2019b). To analyse the spiking events, the low-pass filtered raw traces were further low-pass filtered at 10 Hz to remove subthreshold events and baseline drift. The optical traces were then temporally binned at 1 kHz. Spike detection was carried out using the “get_spikes” function in the Voltage Imaging pipeline (https://github.com/ahrens-lab/VoltageImaging_pipeline^1^ Three parameters (rolling window size, threshold sets for spike size and standard deviation) were optimized by comparing with the ground truth electrophysiology data to achieve minimal false positive spikes while preserving as many true positive spikes as possible. Peristimulus time histogram (PSTH, George Gerstein, U. of Pennsylvania, Neuroscience) was calculated in Python (Python 3.8) with optimized window sizes and bin sizes to show millisecond-level correlation (50 ms window, 2 ms bin) and second-level correlation (1 s window, 100 ms bin) separately^29^. To compare with traces that have the same firing rate, 100 randomly shuffled cell pairs were generated by shuffling spike times of the original two traces. Cross-correlation of subthreshold activities were analysed by normalizing the activity traces and using the “spicy.signal.correlate” function in Python with full lags. 100 randomly shuffled cell pairs were generated by pairing 10-seconds recording traces from different cells in different brain slices.

## Supplementary data

**Supplementary Table 1.** Image sensor specification value comparison The photo-response non uniformity (PRNU) is the spatial non-uniformity in sensitivity between pixels. The dark noise response non-uniformity (DRNU) is the spatial non-uniformity in noise between pixels. N/A, not applicable; PDP, photon detection probability (does not include fill factor). *The incident light is digitised by converting photons to electrons (e^-^). Full well capacity is the number of electrons that can be stored within the well. Table adapted from ^16^.

| Type | emCCD | sCMOS |  | Binary SPAD |
| --- | --- | --- | --- | --- |
| Model | Andor iXon Ultra 888 | Zyla 4.2 PLUS | Hamamatsu ORCA-Flash4.0 V2/V3 | SPCImager |
| GEVIs | QuasAr1&2<sup>23</sup><br>SomArchon<sup>24</sup><br>ASAP3<sup>2</sup> | SomArchon<sup>23</sup><br>Voltron-JF<sub>525</sub>-HTL<sub>1</sub> | QuasAr1&2<sup>23</sup><br>paQuasAr3<sup>25</sup><br>Voltron-JF<sub>525</sub>-HTL<sub>1</sub><br>Positron<sup>20</sup> | Voltron-JF<sub>525</sub>-HTL<br>(this manuscript) |
| Full-well capacity (e<sup>-</sup>)<sup>*</sup> | 80,000 | 30,000 | 30,000 | 1 |
| Dynamic range | 80,000:1 | 33,000:1 | 37,000:1 | 100,000:1 |
| Peak Quantum efficiency | >95% (500-600nm) | 82% @ 560nm | 82% @ 560nm | 35% (PDP) @ 450nm |
| Fill factor | 100% | N/A | N/A | 26.8% |
| Array size | 1024 x 1024 | 2048 x 2048 | 2048 x 2048 | 320 x 240 |
| Pixel size (μm) | 13 | 6.5 | 6.5 | 8 |
| Read noise (e<sup>-</sup>) | <1 with EM gain | 0.9 | 1.4 | ~0 |
| Dark noise (dark count rate) | 0.00025 Hz | 0.10 Hz | 0.05 Hz | 25 Hz |
| Non-uniformity | N/A | < 0.1% PRNU | 1% DRNU, 0.5% PRNU | 2% DRNU, 1% PRNU |
| Frame rate | 26 fps (670 fps with 128 x 128 Crop Mode) | 53 fps (26,041 fps from a 1024(h) x 8(v) ROI, 12 bit) | 100 fps (9,329 fps from a 1024(h) x 8(v) ROI, 16 bit) | 9,938.4 fps (binary frames) |
| Power dissipation | 72 W | 25 W (typical) | 55 W | <1 W (sensor only) |

**Supplementary figure 1.**
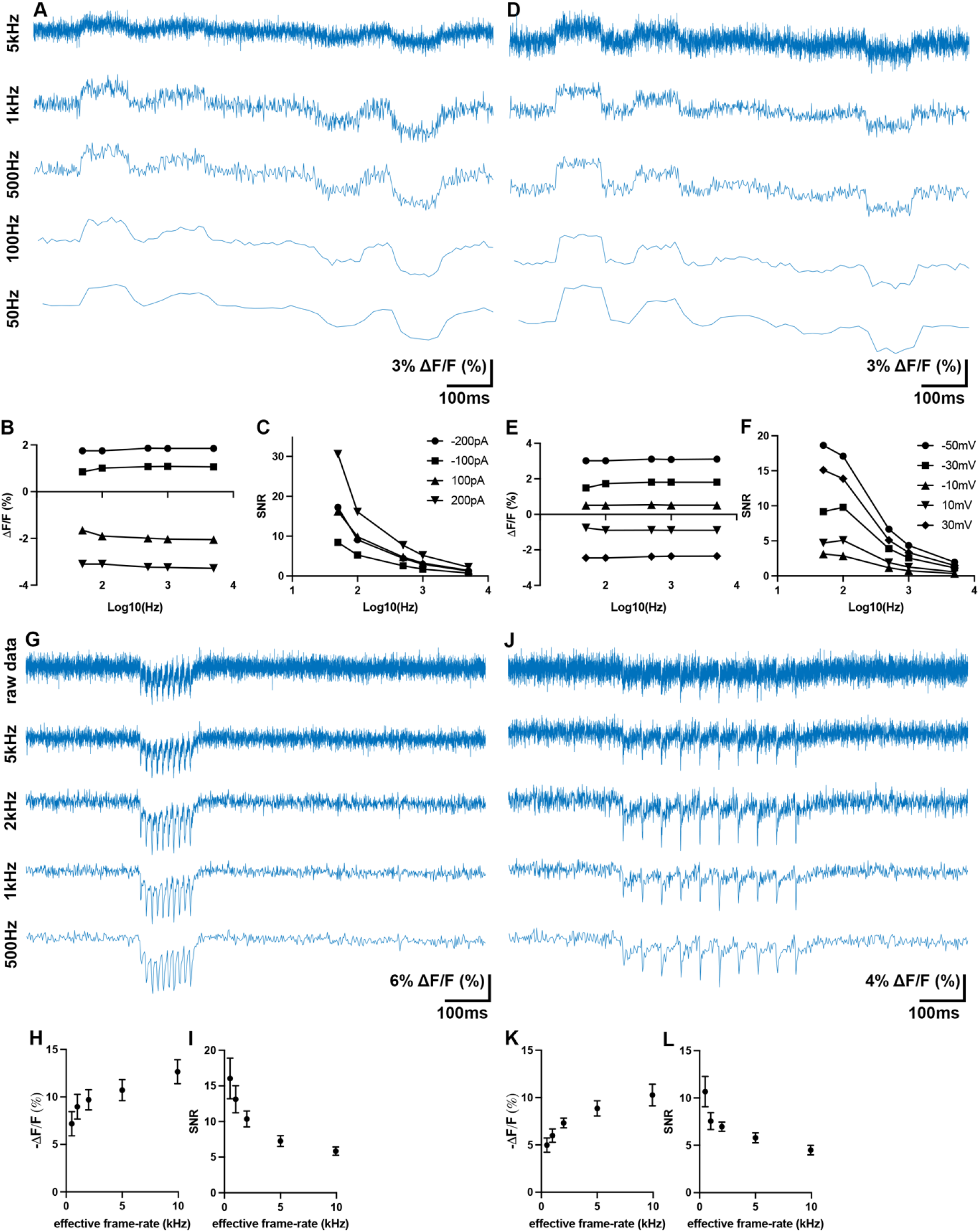
Effects of temporal binning on ΔF/F and SNR. (A, D) Representative optical traces of the same neuron subjected to current step (−200, −100, 100 and 200 pA) (A) and voltage step (−50, −30, −10, 10, 30 mV) (D) stimuli and temporally binned to achieve effective frame-rates of 50 Hz to 5 kHz. (B-C) Relationship between effective frame-rate and ΔF/F (B) and SNR (C) for each of the 4 current step stimuli in (A). (E-F) Relationship between effective frame-rate and ΔF/F (E) and SNR (F) for each of the 4 voltage step stimuli in (D). (G, J) Representative optical traces of the same neuron firing 10 action potentials at 100 Hz (G) and 25Hz (J) were temporally binned to achieve effective frame-rates of 500-5k Hz. (H-I) Relationship between effective frame-rate and averaged ΔF/F (H) and SNR (I) of the 10 spikes in (G). (K-L) Relationship between effective frame-rate and averaged ΔF/F (K) and SNR (L) of the 10 spikes in (J). (H, I, K, L), mean ± SEM.

**Supplementary figure 2.**
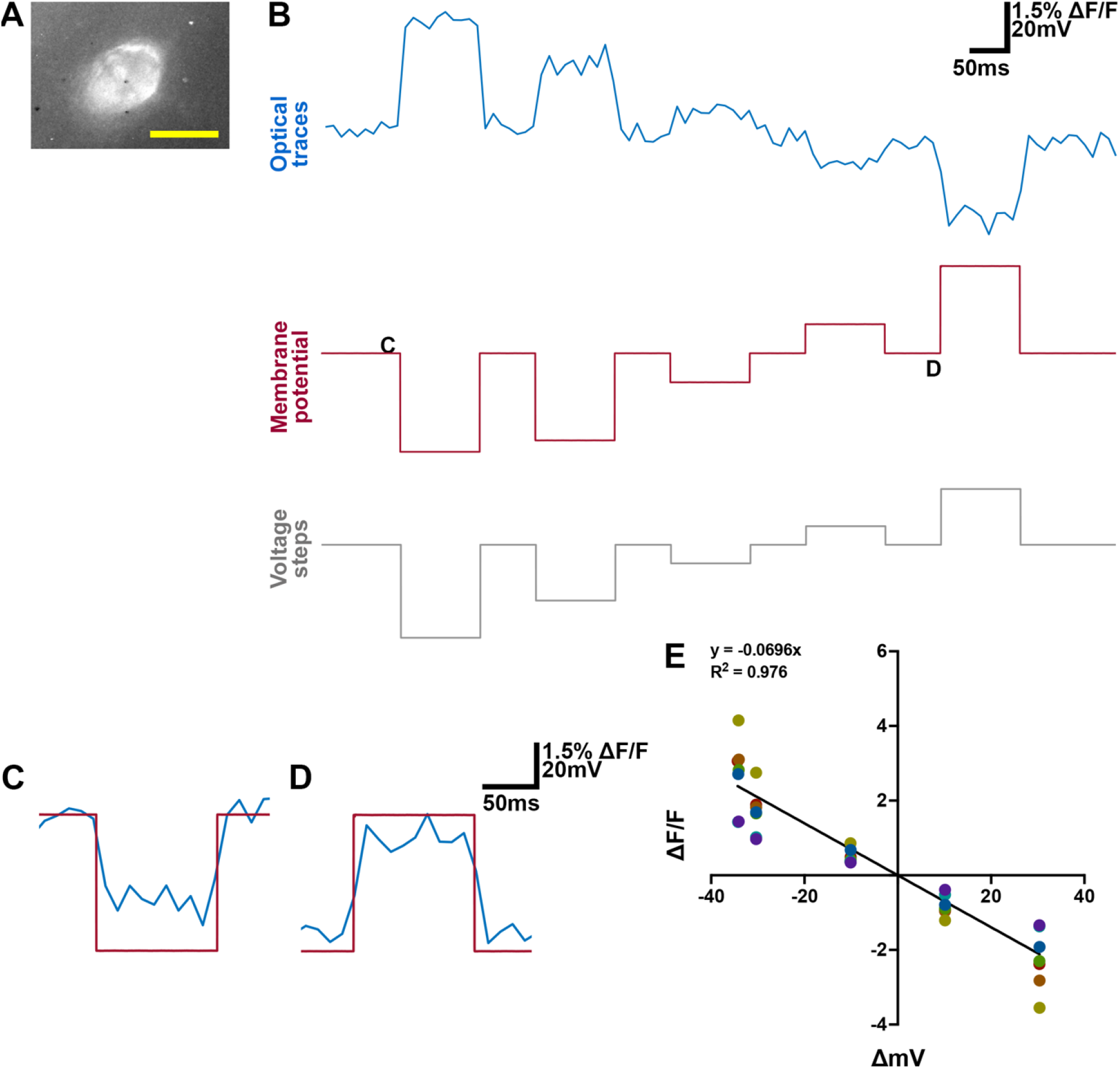
SPAD detection of responses to voltage step stimuli. (A) Image captured by the SPAD array of a patch-clamped neuron expressing Voltron-JF525-HTL, from which the traces in (B-D) was recorded. Scale bar for neuron image: 10μm. (B) Single continuous traces from the neuron in (A) of simultaneous optical (blue) and electrical (red) recordings of membrane potential changes in response to voltage steps(grey). The optical traces were collected at a sampling at 9.9384 kHz, low pass filtered at 2 kHz and temporally binned at 100 Hz. (C-D) Zoom-in of the segments with respective letters of the traces in panel B, with the optical (blue) and electrical (red) traces superimposed. (E) The relationship between ΔF/F and changes in membrane potential (ΔmV) (n = 7 cells). The same cells with different current steps input were marked with the same colour. Simple linear regression was applied with intercepts set at x = 0 and y = 0, p< 0.0001.

**Supplementary figure 3.**
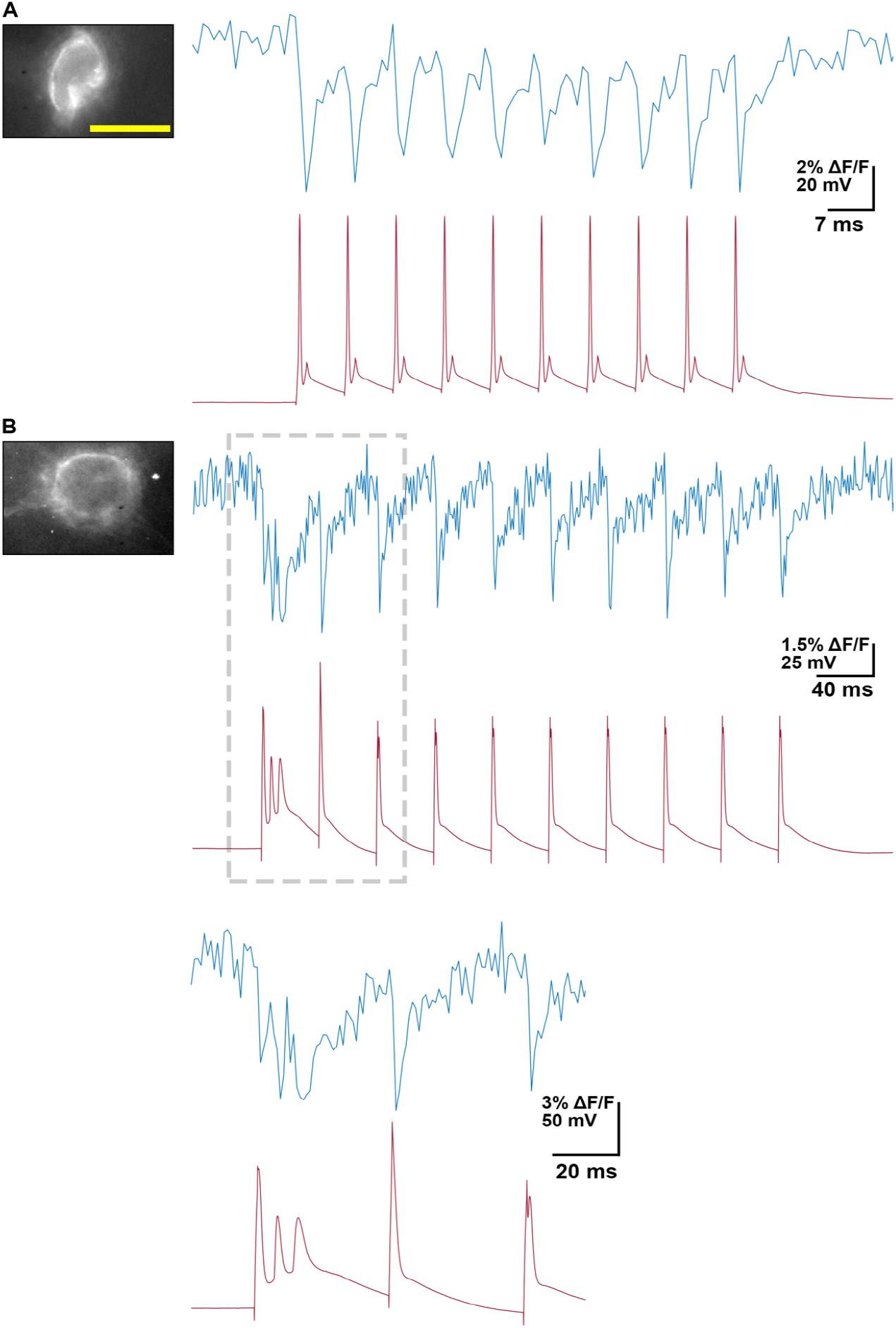
The SPAD image sensor can detect trains of action potential up to 150 Hz and complex spikes reported by Voltron-JF525-HTL. (A) Single continuous traces of simultaneous optical (blue) and electrical (red) recordings of the neuron on the left firing 10 action potentials at a frequency close to 150 Hz. (B) Single continuous traces of simultaneous optical (blue) and electrical (red) recordings of the neuron on the left showing complex spikes fired in response to one current pulse of 2 nA with 1 ms duration, showing SPAD can also record complex spikes reported by Voltron-JF525-HTL. The traces at the bottom show the traces in the segmented rectangle in (B) on an expanded time scale. Scale bar: 10μm.

**Supplementary figure 4.**
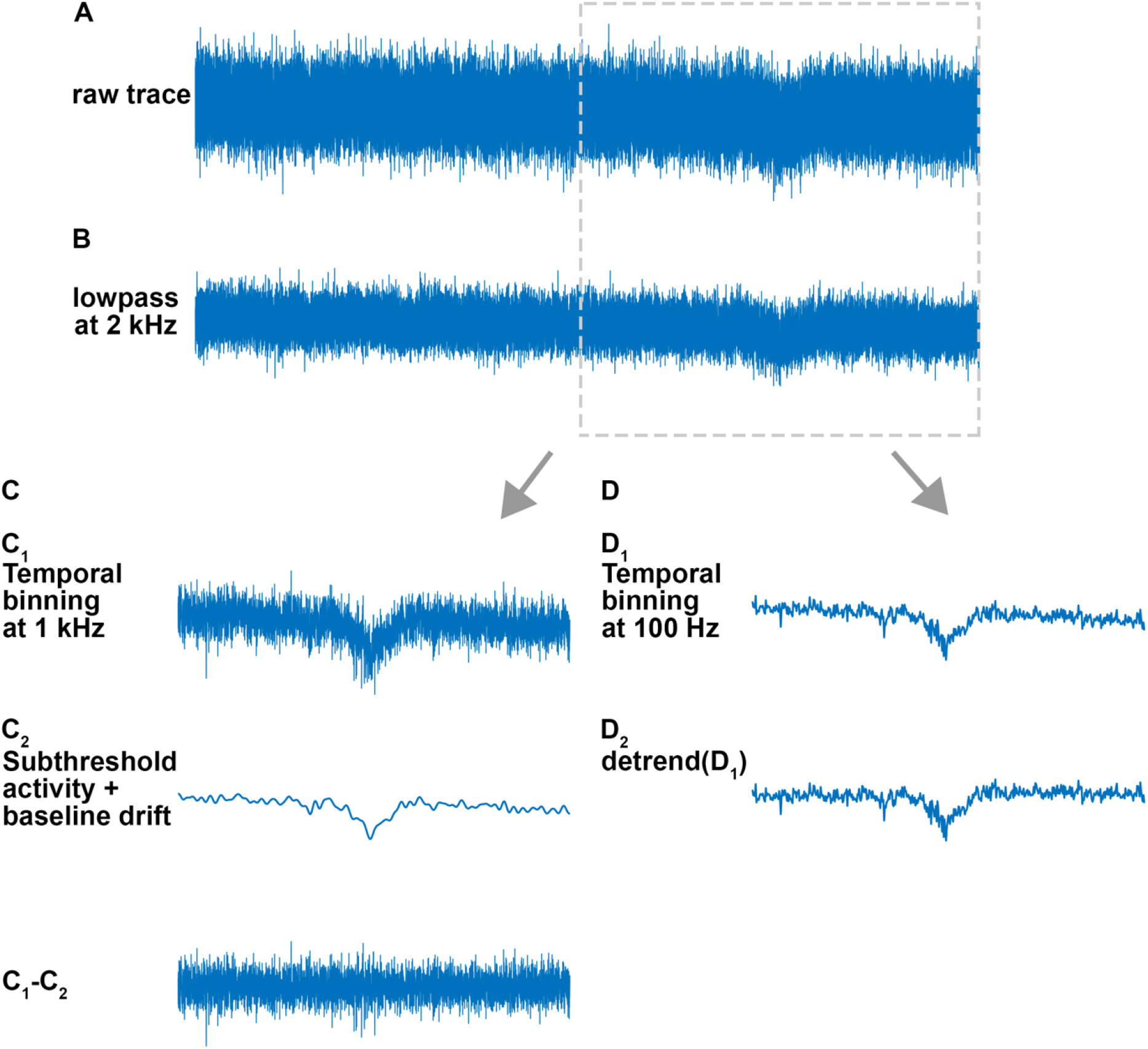
Analysis of sub-threshold and spike components of optical signals. (A-B) Raw 10-second duration optical trace recorded by the SPAD sensor and then low pass filtered at 2 kHz. The region included in the segmented rectangle was used as the example trace for processing steps in (C-D). (C) The optical trace in (B) that was temporally binned to an effective frame rate of 1 kHz (C_1_). Subthreshold activities and baseline-drifting due to photobleaching (C_2_) were removed by low-pass filter at 10 Hz. Spike detection was then carried on the optical traces generated by C_1_-C_2_. (D) Subthreshold activities were revealed by temporally binning the optical traces in (B) at 100 Hz (D_1_) and removing the baseline-drifting due to photobleaching (D_2_).

**Supplementary figure 5.**
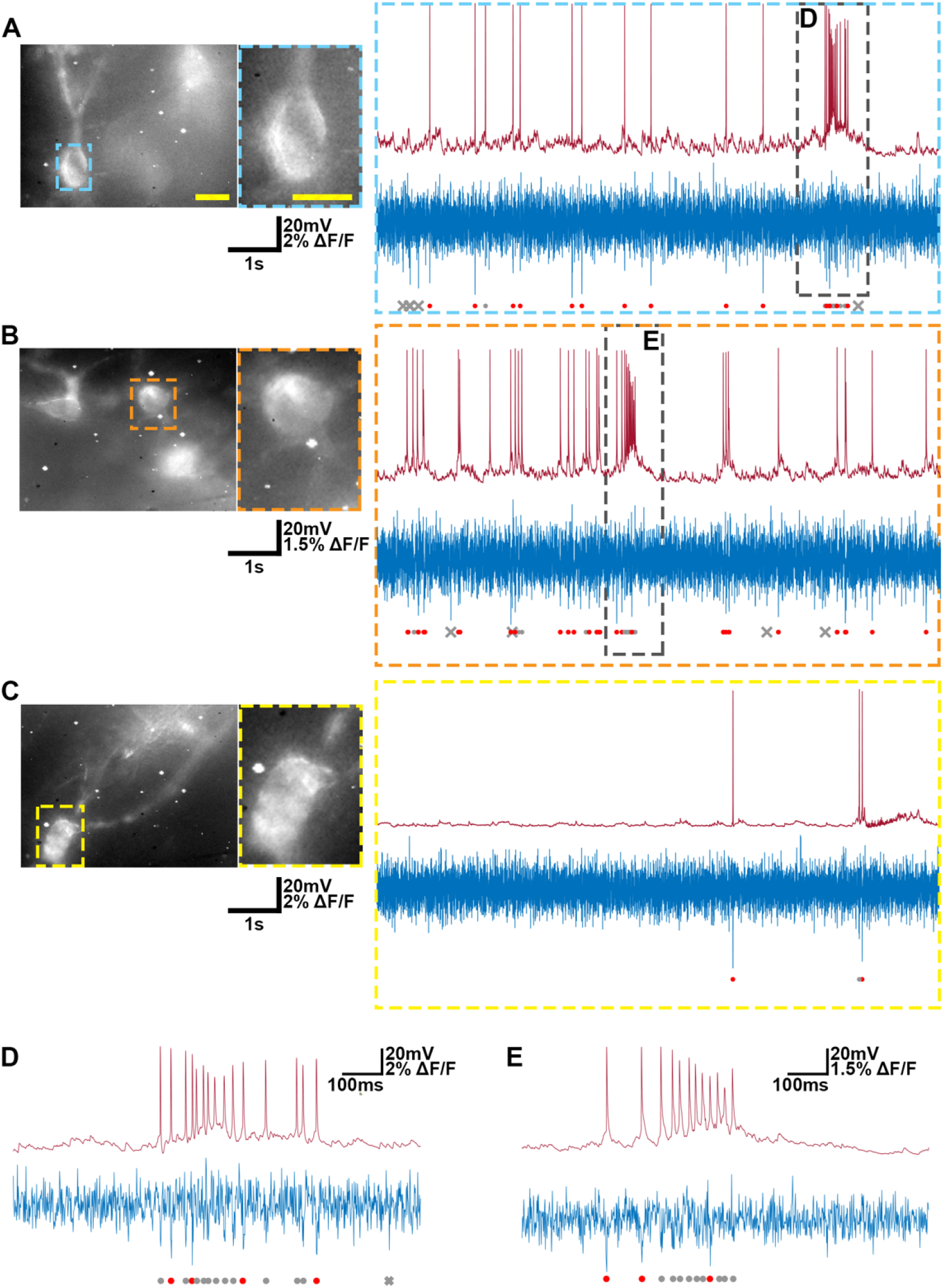
Detection of spikes during epileptiform activity. (A-C) Examples for three neurons of membrane potential recorded with patch-clamp (upper right), the corresponding optical trace baseline-subtracted and binned at 1 kHz as in Supplemental Figure 5 (lower right), and images of the field of view and region of interest for analysis (left). Correctly detected spikes are marked with a red dot, false positive spikes with a grey cross and false negative spikes with a grey dot. Scale bar: 10μm. (D-E) Regions indicated in A and B on an expanded time base to illustrate signal and spike detection during activity bursts.

## Funding, acknowledgements and copyright

We thank Robert Henderson and Ian Duguid for helpful discussions and support. The project was supported by funding from the Wellcome Trust (ISSF3 award IS3-R2.36 to IG and MFN, and Investigator Award 200855/Z/16/Z to MFN), the BBSRC EastBio doctoral training programme, and EPSRC (EP/S001638/1). For the purpose of open access, the author has applied a CC BY public copyright licence to any Author Accepted Manuscript version arising from this submission.

## Data availability

Data will be made available via https://datashare.ed.ac.uk/handle/10283/777.

